# A universal dynamical metabolic model representing mixotrophic growth of *Chlorella* sp

**DOI:** 10.1101/2022.07.12.499674

**Authors:** Bruno Assis Pessi, Caroline Baroukh, Anais Bacquet, Olivier Bernard

## Abstract

An emerging idea is to couple wastewater treatment and biofuel production using microalgae to achieve higher productivities and lower costs. This paper proposes a metabolic modelling of *Chlorella sp*. growing on wastes in mixotrophic conditions, accounting also for the possible inhibitory substrates. A metabolic model considering several possible carbon substrates was developed and run. The addition of several organic carbon substrates such as acetate, butyrate or glucose were tested, along with glycerol, a more realistic substrate from an economical point of view. The metabolic model was built using DRUM framework and consists of 188 reactions and 176 metabolites. After a calibration phase, the model was successfully challenged with data from 122 experiments collected from scientific literature in autotrophic, heterotrophic and mixotrophic conditions. The optimal feeding strategy estimated with the model reduces the time to consume the volatile fatty acids from 16 days to 2 days. The high prediction capability of this model opens new routes for enhancing design and operation in waste valorisation using microalgae.

**Author Summary:** Waste valorisation is one of the current envisaged strategies to make renewable processes more economically advantageous. For example, wastewater treatment can be used to produce biohydrogen from bacteria, through a process called dark fermentation, and to cultivate microalgae for biofuel production. Dark fermentation has, as by-products, organic acids that have inhibitory effects on the growth of microalgae, increasing the time to completely treat the waste. Advances in metabolic knowledge and techniques allow for the deployment of new strategies to improve the efficiency of bioprocesses. In this work, we validate a mathematical model of the metabolism of the microalgae genus *Chlorella* using the DRUM framework for 122 experiments from the scientific literature. This model enables us to apply control and optimisation techniques to provide a strategy to treat wastes coming from dark fermentation processes, overcoming the inhibition of some organic acids. The strategy is able to reduce the time to treat the waste from 16 days to only 2 days. The high prediction capability of this model opens new routes for enhancing design and operation in waste valorisation using microalgae.

## 1. Introduction

Microalgae have been extensively studied during the past decade. Indeed, certain species are capable of producing lipids or carbohydrates that can in turn be converted into biofuel (Wijffels and Barbosa, 2010). Microalgae use light energy, via photosynthesis, to fix carbon dioxide. Not only, their growth rate is much faster than that of higher plants, but they can also be cultivated in wastewater. An emerging idea suggests combining wastewater treatment with the production of biofuels to minimize its costs (Christenson and Sims, 2011). However, even though the production efficiency appears to be attractive, many optimization steps still need to be carried out for this process to become sufficiently cost-effective and environmentally-friendly (Tan et al., 2018).

The growth of microalgae on primary effluents or effluents from a first fermentation phase such as dark fermentation has already been studied (Baroukh et al., 2017, Turon et al., 2015b,a). The advantage of dark fermentation is that complex and non-assimilable compounds can be transformed into usable carbon sources for microalgae. In particular, thanks to these fermentation processes, which are achieved by anaerobic bacteria and Archaea, waste and effluents can be converted into Volatile Fatty Acids (VFAs). VFAs could thus represent a carbon source for microalgae, while microalgae get nitrogen and phosphorus from ammonium and phosphate in the wastewater. The VFA mixture resulting from dark fermentation typically comprises about 30% acetate, 70% butyrate, and sometimes lactate (Rafrafi et al., 2013). Nevertheless, a major drawback is the presence of butyrate in these effluents, what exerts a strong inhibition on algal growth (Hu et al., 2012, Turon et al., 2015b).

In order to eliminate the inhibition effect of butyrate and thus to accelerate growth in dark fermentation effluents, the addition of another carbon substrate to the medium could be considered. Indeed, these substrates could enhance biomass production, until reaching the necessary critical threshold to overcome the inhibition. Glucose could ideally play this role, although it is unattractive due to its cost. Acetate could be a possible alternative as it is less costly; however it generates large variations in pH. Glycerol is a by-product of biodiesel synthesis by transesterification. Therefore, its addition would not increase the production cost of the process very significantly.

The objective of this article was to develop a universal multi-substrate reduced metabolic model of the growth of *Chlorella sp*.. The reduced metabolic model allowed to provide accurate phenotype predictions, while enabling the use of computationally demanding analysis frameworks (Singh and Lercher, 2020). This model was key to determine the optimal strategy for substrate addition, in fermentation effluents to maximize productivity. *Chlorella sp*. was selected as it demonstrated its potential in associating biofuel production with effluent treatment Casagli et al. (2021b). Indeed, this species can accumulate up to 50% of its dry weight in lipids, essentially in the form of triacylglycerol (TAG). In addition, it has the capacity to grow in dark fermentation effluents, under heterotrophic or mixotrophic conditions (Turon et al., 2015a). The model developed here represents, in detail, the growth conditions, under different autotrophic, heterotrophic and mixotrophic conditions and for various substrates. The model was validated in different cultivation conditions using the abundant literature available on autotrophic, heterotrophic or mixotrophic growth of *Chlorella*. To this end, data from 122 experiments was extracted from 15 publications, amounting to more than 2600 concentration data points (see Table 3).

## 2. Materials and methods

### 2.1. General Principles of the DRUM approach

To develop the model, we used the DRUM (Dynamic Reduction of Unbalanced Metabolism) approach. The full description and complete explanation of the approach is, available in Baroukh et al. Baroukh et al. (2014).

Briefly, the metabolism of a microorganism can be described by its metabolic network composed of a set of *n*_*r*_ biochemical reactions (here *n*_*r*_ = 188) involving *n*_*m*_ metabolites (here *n*_*m*_ = 176) and represented by the stoichiometric matrix 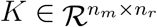. The biomass B is produced from a set of substrates S and excretes a set of products P. In a perfectly mixed reactor with a constant volume, the system can be described by the following set of ordinary differential equations:

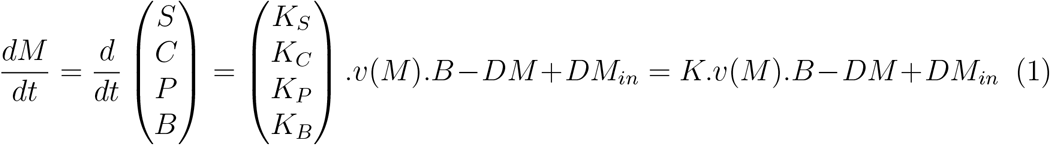

where M represents the vector of the concentrations of metabolites composed of substrate (S), intracellular metabolites (C), excreted products (P) and biomass (B). *M*_*in*_ is the influent concentration of these quantities. The dilution rate of the reactor (ratio of influent flow rate over the reactor volume) is *D* (*D* = 0 for a batch process).

All the concentrations are expressed as total concentrations in the solution. 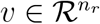 is the reaction kinetic vector, while the matrices 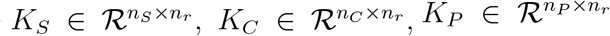 and 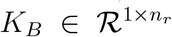 correspond, respectively, to the stoichiometric matrices of substrates S, products P, intracellular metabolites C and biomass B (*n*_*S*_ + *n*_*C*_ + *n*_*P*_ + 1 = *n*_*m*_).

In most of metabolic models, intracellular metabolites are generally assumed to be quasi-stationary 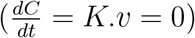, i.e. they are assumed to be consumed as soon as they have been synthesised. However, in the case of microalgae, this hypothesis has proven to be false for certain of its metabolites (marked as A) during mixotrophic or autotrophic growth (Baroukh et al., 2014, 2017). The DRUM method (Baroukh et al., 2014), consists in dividing the metabolic network into n quasi-stationary subnetworks (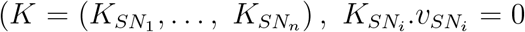 *for i* ∈ 1, …, *n*). These are linked by *A* metabolites that are, in contrast, non-stationary and can accumulate and be later consumed. This division into subnetworks is justified by the presence of metabolic pathways that correspond to metabolic functions, to reaction groups that are regulated simultaneously and to the presence of compartments within the cell. Cellular mechanisms are therefore employed for assessing the subnetwork. Hence, *A* metabolites can either be found at the junction of several metabolic pathways, or they can be transported from one compartment to another, or they can be final products that accumulate in the cell. The system of ordinary differential equations (1) therefore becomes:

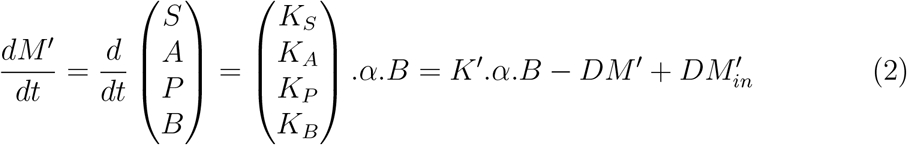

With *K*^′^ the stoichiometric matrix of macroscopic reactions obtained through the analysis of elementary modes (Schuster et al., 1999) on the subnetworks, and *α* the kinetic vector associated to these macroscopic reactions. B now represents the structural biomass, i.e. the fraction of biomass that does not contain the inert compartments of reserve A. The total biomass can be deduced using a mass balance of the elemental compounds (C, N, P, O, …).

### 2.2. Construction of the model

The core of the metabolic network from Baroukh et al. (2017) has been used and modified in order to add the glucose and glycerol consumption pathways (Figure 1). This network contains the central autotrophic, mixotrophic and heterotrophic metabolic pathways including photosynthesis, glycolysis, the pentose phosphate pathway, the Krebs cycle, oxidative phosphorylation and the synthesis of chlorophyll, carbohydrates (e.g. starch), amino-acids and nucleotides. The synthesis pathways of macromolecules such as proteins, lipids, starch, DNA, RNA as well as the functional biomass are represented through macroscopic reactions.

**Figure 1:**
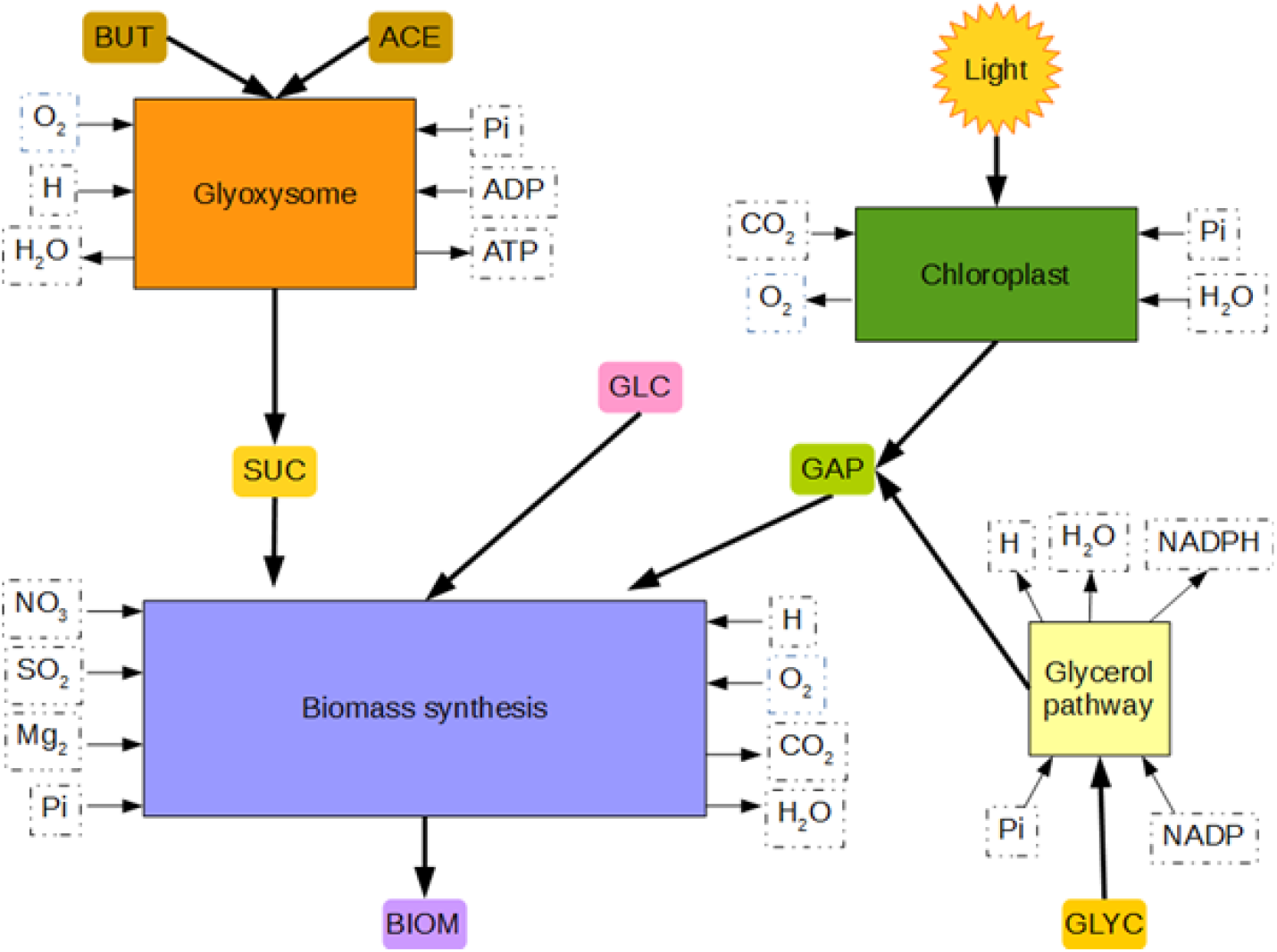
Considered metabolic subnetworks to represent growth of *Chlorella* on a mixture of glycerol, glucose, acetate and butyrate

The DRUM method requires the partitioning of the metabolism into subnetworks as well as the identification of the metabolites, in between the subnetworks, which can accumulate. The subnetworks are defined by their metabolic function and/or their affiliation to a cellular compartment. Different partitions among the 188 reactions have been tested, with a view to select the one which minimized the number of parameters to assess while providing a correct representation of the experimental data. The network is divided into four subnetworks (Figure 1), corresponding to, 1) the glyoxysome, 2) the chloroplast, 3) the absorption of glycerol and 4) the synthesis of biomass. The glyoxysome and chloroplast subnetworks remain unchanged in comparison with the initial Baroukh et al. (2017) model.

The macroscopic reactions associated to each subnetwork are deduced using the method for elementary mode analysis (Klamt and Stelling, 2003). The Matlab “efmtool” has been run to calculate the Elementary Flux Modes (EFMs) (Terzer and Stelling, 2008). In total, 86, 142 EFMs including 3, 310 associated to futile cycles (dissipation of carbon substrate in the form of CO2) have been achieved. These macroscopic reactions aim at determining the mass fluxes in the different parts of the network by assembling reactions belonging to the same kinetics.

### 2.3. Analysis of the sub-networks

#### 2.3.1. Motivations

In this section, we present the macroscopic reactions which result from the reduction of each subnetwork by the computation of the EFMs. As recommended by Baroukh et al. (2014), the kinetics must be chosen using minimal hypotheses, and when possible applying a mass action law. A list of all sub-networks and the macroscopic reactions can be found at Table 1.

**Table 1:** List of macroscopic reactions and the respective subnetwork. For biomass production the stoichiometric values may differ if the nitrogen source is nitrate or ammonium.

|  | Subnetwork | Macroscopic reaction |
| --- | --- | --- |
| MR<sub>1</sub> | Glyoxysome | $2 \text{ ACE} + 3.5 \text{ H} + 0.5 \text{ O}_2 \rightarrow \text{SUC} + 0.5 \text{ H}_2\text{O}$ |
| MR<sub>2</sub> | | $\text{BUT} + 7 \text{ H} + 1.5 \text{ O}_2 \rightarrow \text{SUC} + 5 \text{ H}_2\text{O}$ |
| MR<sub>3</sub> | Chloroplast | $\text{Light} + 3 \text{ CO}_2 + 2 \text{ H}_2\text{O} + \text{Pi} \rightarrow \text{GAP} + 3 \text{ O}_2$ |
| MR<sub>4</sub> | Glycerol pathway | $\text{GLYC} + \text{Pi} \rightarrow \text{GAP} + \text{H}_2\text{O}$ |
| MR<sub>5</sub>(NH<sub>4</sub>) | Biomass synthesis | $4.15 \text{ GAP} + 2.54 \text{ O}_2 + 0.99 \text{ NH}_4 + 0.02 \text{ SO}_4 + 0.01 \text{ Mg}_2 \rightarrow$<br>$\text{B} + 0.99 \text{ H} + 2.90 \text{ H}_2\text{O} + 3.92 \text{ CO}_2 + 4.02 \text{ Pi}$ |
| MR<sub>5</sub>(NO<sub>3</sub>) | | $4.64 \text{ GAP} + 2.04 \text{ O}_2 + 0.99 \text{ NO}_3 + 0.98 \text{ H} + 0.02 \text{ SO}_4 +$<br>$0.01 \text{ Mg}_2 \rightarrow \text{B} + 5.39 \text{ CO}_2 + 2.90 \text{ H}_2\text{O} + 4.51 \text{ Pi}$ |
| MR<sub>6</sub>(NH<sub>4</sub>) | Biomass synthesis | $4.15 \text{ SUC} + 7.30 \text{ H} + 4.61 \text{ O}_2 + 0.99 \text{ NH}_4 + 0.12 \text{ Pi} +$<br>$0.02 \text{ SO}_4 + 0.01 \text{ Mg}_2 \rightarrow \text{B} + 7.04 \text{ H}_2\text{O} + 8.06 \text{ CO}_2$ |
| MR<sub>6</sub>(NO<sub>3</sub>) | | $4.90 \text{ SUC} + 5.28 \text{ O}_2 + 0.99 \text{ NO}_3 + 0.12 \text{ Pi} + 10.78 \text{ H} +$<br>$0.02 \text{ SO}_4 + 0.01 \text{ Mg}_2 \rightarrow \text{B} + 11.07 \text{ CO}_2 + 8.31 \text{ H}_2\text{O}$ |
| MR<sub>7</sub>(NH<sub>4</sub>) | Biomass synthesis | $2.07 \text{ GLC} + 2.54 \text{ O}_2 + 0.99 \text{ NH}_4 + 0.12 \text{ Pi} + 0.02 \text{ SO}_4 +$<br>$0.01 \text{ Mg}_2 \rightarrow \text{B} + 3.91 \text{ CO}_2 + 7.04 \text{ H}_2\text{O} + 0.99 \text{ H}$ |
| MR<sub>7</sub>(NO<sub>3</sub>) | | $2.34 \text{ GLC} + 2.14 \text{ O}_2 + 0.99 \text{ NO}_3 + 0.12 \text{ Pi} + 0.98 \text{ H} +$<br>$0.02 \text{ SO}_4 + 0.01 \text{ Mg}_2 \rightarrow \text{B} + 5.49 \text{ CO}_2 + 7.63 \text{ H}_2\text{O}$ |

**Table 2:**
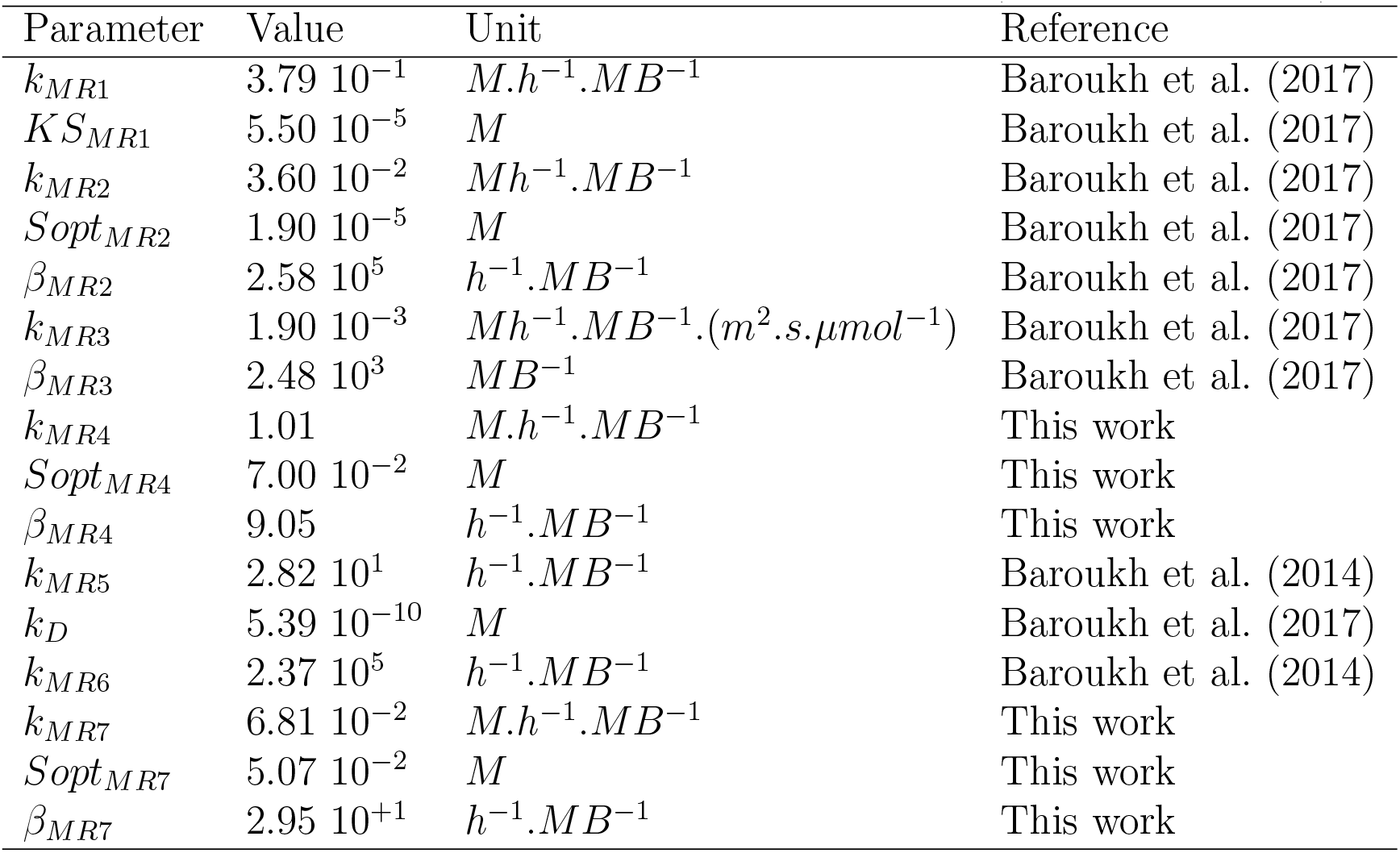
Kinetic parameters obtained after model calibration (MB: mole of biomass)

#### 2.3.2. Glyoxysome subnetwork

The glyoxysome pathway consists of 26 reactions, from which 8 are exchange reactions. The glyoxysome is the peroxysome compartment where the glyoxylate cycle occurs. Here carbon compounds are converted to succinate, also allowing the production of glucose from lipids. In this compartment, fatty acids can be used as a source of energy and carbon for growth when no photosynthesis takes place. Two EFMs have been achieved for this subnetwork (MR1 and MR2). In the glyoxysome, butyrate and acetate are converted into acetyl-CoA, which is in turn converted, via the glyoxylate cycle, into succinate. The succinate then enters the cytosol and is injected into the Krebs cycle, thus producing the different metabolites necessary for the synthesis of biomass.

Butyrate induces an inhibition on algal growth under heterotrophic and mixotrophic conditions (Turon et al., 2015a). Furthermore, acetate inhibits the absorption of butyrate, thus leading to diauxic growth (Turon et al., 2015a). Thereby, Michaelis-Menten kinetics have been proposed to describe the absorption of acetate (*α*_*MR*1_). For butyrate (*α*_*MR*2_) Haldane kinetics have been chosen with an inhibition by acetate term.

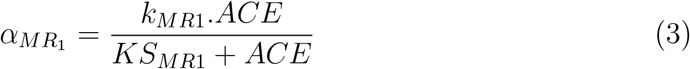

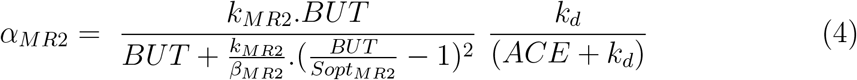

#### 2.3.3. Chloroplast subnetwork

The chloroplast subnetwork is composed of 21 reactions, from which 7 are exchange reactions. The glycerate-3-phosphate produced by photosynthesis is assumed to be transferred from the chloroplast towards the cytosol where it can be converted by glycolysis into glucose-6-phosphate or pyruvate. These metabolites are essential for the synthesis of functional biomass.

When autotrophic growth occurs, light is assumed to be the main limitation for this reaction. When algae are growing on a turbid medium like wastewater, light intensity stays low and the photoinhibition mechanisms can be ignored Martínez et al. (2018). Thus, photosynthesis rate is assumed to be linearly depending upon the average light intensity *I*_*μ*_ in the culture (see Equation 6).

Moreover, light attenuation in the culture medium is expected to follow the Beer-Lambert law. The light intensity at depth *z* depends on the incident light *I*_0_ and the extinction coefficient *α* due to the biomass (the turbidity of the medium without algae is negligible):

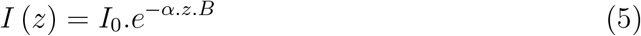

Hence, the average light intensity of the culture medium in the reactor of depth *L* is given as follows (with *β*_*MR*3_ = *α*.*L*):

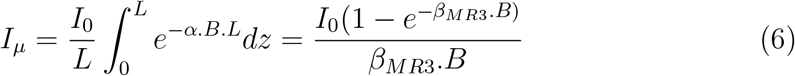

Hence, the kinetics in the chloroplast subnetwork is given by:

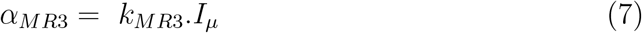

#### 2.3.4. Glycerol absorption subnetwork

The glycerol pathway subnetwork consists of 5 core reactions, plus the exchange reactions. Only one EFM was found for the glycerol absorption subnetwork (MR4).

The glycerol present in the medium is transferred to the cytosol. Within three steps, it is then transformed into glycerate-3-phosphate. During glycolysis, this glycerate-3-phosphate is then used for the synthesis of precursor metabolites that are in turn required for the synthesis of functional biomass. Reaction kinetics with inhibition were assumed for glycerol assimilation (*α*_*MR*4_), as inhibition can be observed in the literature (Chen and Walker, 2011, Ma et al., 2016, Liang et al., 2009).

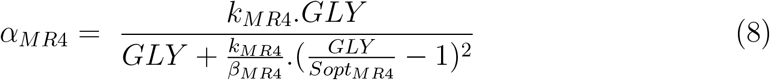

#### 2.3.5. Functional biomass synthesis subnetwork

The reactions for the synthesis of lipids, proteins, DNA, RNA, chlorophyll and carbohydrates are all lumped together in the functional biomass synthesis subnetwork. This subnetwork includes glycolysis, the Krebs cycle, oxidative phosphorylation, the pentose phosphate pathway, carbohydrate, lipid, amino-acid and nucleotide synthesis, as well as the assimilation of nitrogen, sulphur and glucose. In total there are 141 reactions in the functional biomass subnetwork.

This subnetwork generated 86, 167 EFMs, including 3310 that did not produce biomass. Nearly all of the calculated EFMs are part of the biomass synthesis network. They can be sorted by using a similar method to the FBA (Flux Balance Analysis). The cell is presumed to maximise its biomass growth on each substrate (Orth et al., 2010), thus minimising the loss of carbon as CO2. Therefore, for each substrate, the EFM presenting the highest GAP/BIOM, SUC/BIOM and GLC/BIOM yields were selected. In this way, the use of GAP, SUC or GLC for the synthesis of biomass can be described thanks to three macroscopic reactions (MR5, MR6 and MR7). The yield of biomass on the carbon substrate is different i f t he nitrogen source is *NH*_4_ or *NO*_3_, in Table 3 we show the macroscopic reactions for both cases.

**Table 3:** Experiments for each substrate and their references. In parenthesis the number of experiments in mixotrophic conditions

| Substrate | # exp. | Species | References |
| --- | --- | --- | --- |
| Glycerol | 12 (9) | *C. sorokiniana* | León-Vaz et al. (2019) |
|  |  | *C. sp.* | Sen and Martin (2018) |
|  |  | *C. protothecoides* | Chen and Walker (2011), O’Grady and Morgan (2011) |
|  |  | *C. vulgaris* | Ma et al. (2016) |
| Glucose | 22 (11) | *C. sorokiniana* | León-Vaz et al. (2019), Li et al. (2013, 2014) |
|  |  | *C. protothecoides* | Espinosa-Gonzalez et al. (2014), Chen and Walker (2011) |
|  |  | *C. pyrenoidosa* | Ogbonna et al. (1997) |
| Glucose/Glycerol | 2 (2) | *C. protothecoides* | O’Grady and Morgan (2011) |
| Acetate | 40 (25) | *C. sorokiniana* | León-Vaz et al. (2019), Turon et al. (2015a,b), Chen et al. (2017a,b), Xie et al. (2020a) |
|  |  | *C. saccharophila* | Xie et al. (2020b) |
| Butyrate | 10 (4) | *C. sorokiniana* | Turon et al. (2015b,a) |
| Acetate/Butyrate | 23 (6) | *C. sorokiniana* | Turon et al. (2015b,a) |
| Autotrophic | 13 | *C. sorokiniana* | Li et al. (2014), León-Vaz et al. (2019), Turon et al. (2015b,a) |
|  |  | *C. sp.* | Sen and Martin (2018) |

Glycerate-3-phosphate originates from the chloroplast and from the assimilation of glycerol. It is injected into the glycolysis so as to produce the necessary metabolites for growth (*α*_*MR*5_). MR5:

In the functional biomass synthesis subnetwork, the kinetics are supposed to be linear with respect to glycerate-3-phosphate (*GAP*) and succinate (*SUC*):

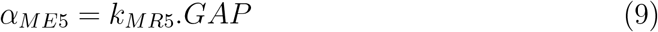

Succinate originates from the glyoxysome. It enters the Krebs cycle, thus also leading to the production of metabolites required for growth (*α*_*MR*6_). MR6:

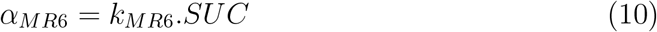

Glucose added to the medium is transferred to the cytosol where it enables the production of biomass. Glucose consumption is assumed to follow a substrate inhibition kinetics (*α*_*MR*7_) (Azma et al., 2011, Wu and Shi, 2007, Liang et al., 2009). MR7:

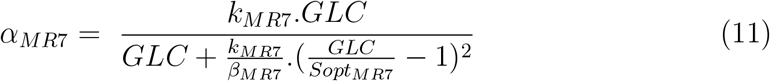

### 2.4. Global dynamics of the network based on the elements that can accumulate

Finally, the dynamical evolution of the metabolic fluxes associated to the 188 reactions involved can be derived from a system with 17 ordinary differential equations comprising 17 metabolites and 7 macroscopic reactions:

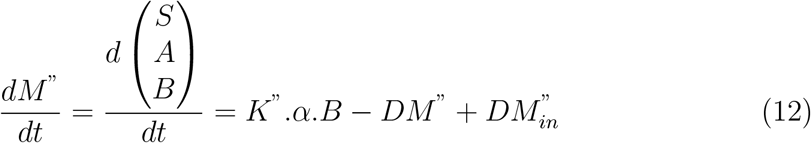

Where *M* ^”^ is the metabolite vector (17×1) comprising the substrates S, the metabolites that can accumulate A (SUC and GAP) and the functional biomass B. K’ is the stoichiometric matrix (17×7) of the macroscopic reactions and *α* the associated kinetics vector (7×1). Moreover, the total biomass comprising the functional biomass and the metabolites A can be described as follows:

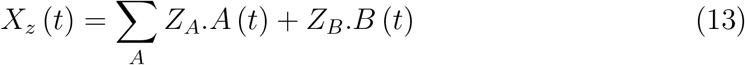

Where Z is a chemical element (*Z* ∈ {*C*; *N* ; *O*; *H*; *P* ; …}), *Z*_*A*_ and *Z*_*B*_ are the number of chemical elements Z per mole of metabolites A and biomass B, *A*(*t*) and *B*(*t*) are the concentrations of A and B at time t *X*_*Z*_(*t*) is the concentration of the chemical element in the total biomass X at time t.

Finally, the metabolic fluxes within the whole network can be derived from the *α* kinetics and the elementary modes associated to the 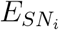, *i* ∈ 1, 2, 3 subnetworks:

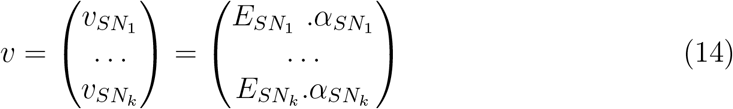

### 2.5. Reduced model calibration

In order to calibrate, and then validate the model, various experiments described in the literature have been used. In total the 122 selected experiments (see Table 3) gather data on growth i) under autotrophic conditions, without any organic carbon input and submitted to light intensities ranging from 30 to 540 E.m-2.s-1 ii) under heterotrophic conditions, without any light, and with varying concentrations in acetate, butyrate, glucose and glycerol, pure or combined iii) under mixotrophic conditions, with light and varying concentrations in acetate, butyrate, glucose and glycerol. Depending on the studies, different combinations of these substrates were considered.

Only the kinetic macro parameters for glucose and glycerol consumption were calibrated, the rest of the kinetic parameters for the macro reactions are taken from Turon et al. (2015b). The calibration was done following a two-step process. First a stochastic global optimizer, Differential Evolution algorithm (Storn and Price, 1997), calculates the set of parameters minimizing the relative error between model and experimental data of biomass and substrate concentration over time. This parameter set is, then, used as initial point in a Markov Chain Monte Carlo sampler, which returns the parameters set inside a confidence interval (Foreman-Mackey et al., 2013). Glucose kinetics parameters were calibrated using concentration data from 5 experiments of Li et al. (2013), while glycerol parameters were fitted using data from 6 experiments of Ma et al. (2016).

## 3. Results

### 3.1. Validation of the model

The experimental data not used during the calibration stage were used to validate the model. A coherent set of experiments, representing various experimental conditions, was kept for this validation stage (see Table 3). We used a Monte Carlo method to select parameters (Foreman-Mackey et al., 2013). We considered a ±20% uncertainty in the initial concentration of substrates. In Figures 2 and 5, the results of the model simulations are compared with the experimental data. The line represents the average of 100 simulations and the colored region represents ±1 standard deviation. As illustrated in Figure 2 for single substrates, the model is capable of suitably predicting the production of biomass and the consumption of substrates, whether in autotrophic, heterotrophic or in mixotrophic conditions. Furthermore, the model is still accurate when there are two substrates (Figure 5). More generally, the predictive performance of the model is summarized in the Taylor diagram in Figure 3 for the whole data set. This diagram represents at the same time the standard deviation of the biomass prediction error and the Pearson correlation coefficient (Taylor, 2001). It illustrates both the centered and reduced quadratic errors between the experimental data sets and the associated simulations, as well as the correlation between the model and the data. It thus summarizes the degree of resemblance between the data and the simulations for the vast range of considered data. Indeed, the closer a data point is to (0;1), the better the model reproduces the experimental data (Taylor, 2001). Figure 4 represents all the data points versus model prediction, also demonstrating the goodness of fit of the model.

**Figure 2:**
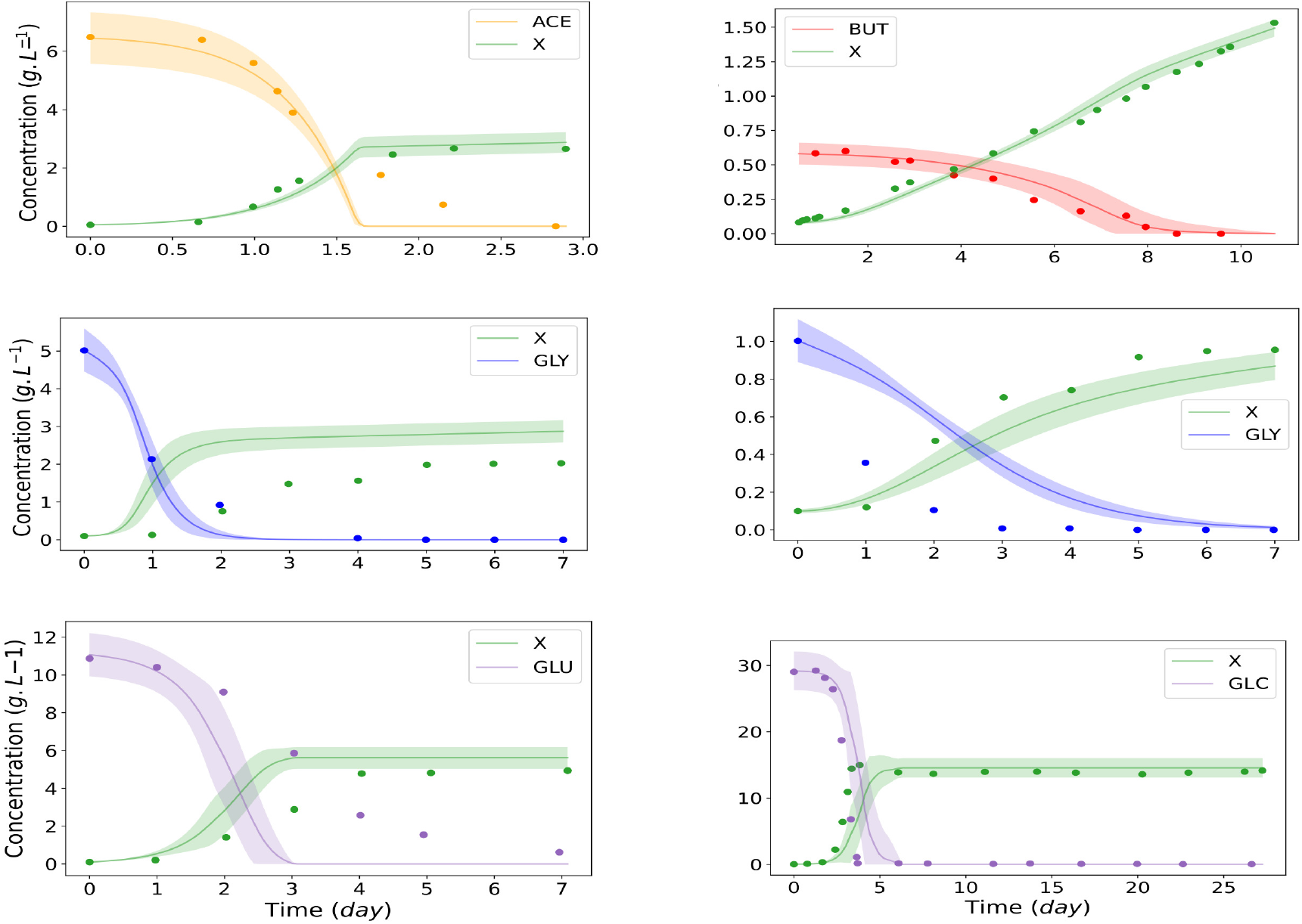
Model validation for various conditions of heterotrophic and mixotrophic growth. Top: Acetate and butyrate in mixotrophic conditions. Middle: Glycerol in heterotrophic conditions. Down: Glucose in heterotrophic growth. Colored regions represent ±1 standard deviation.

**Figure 3:**
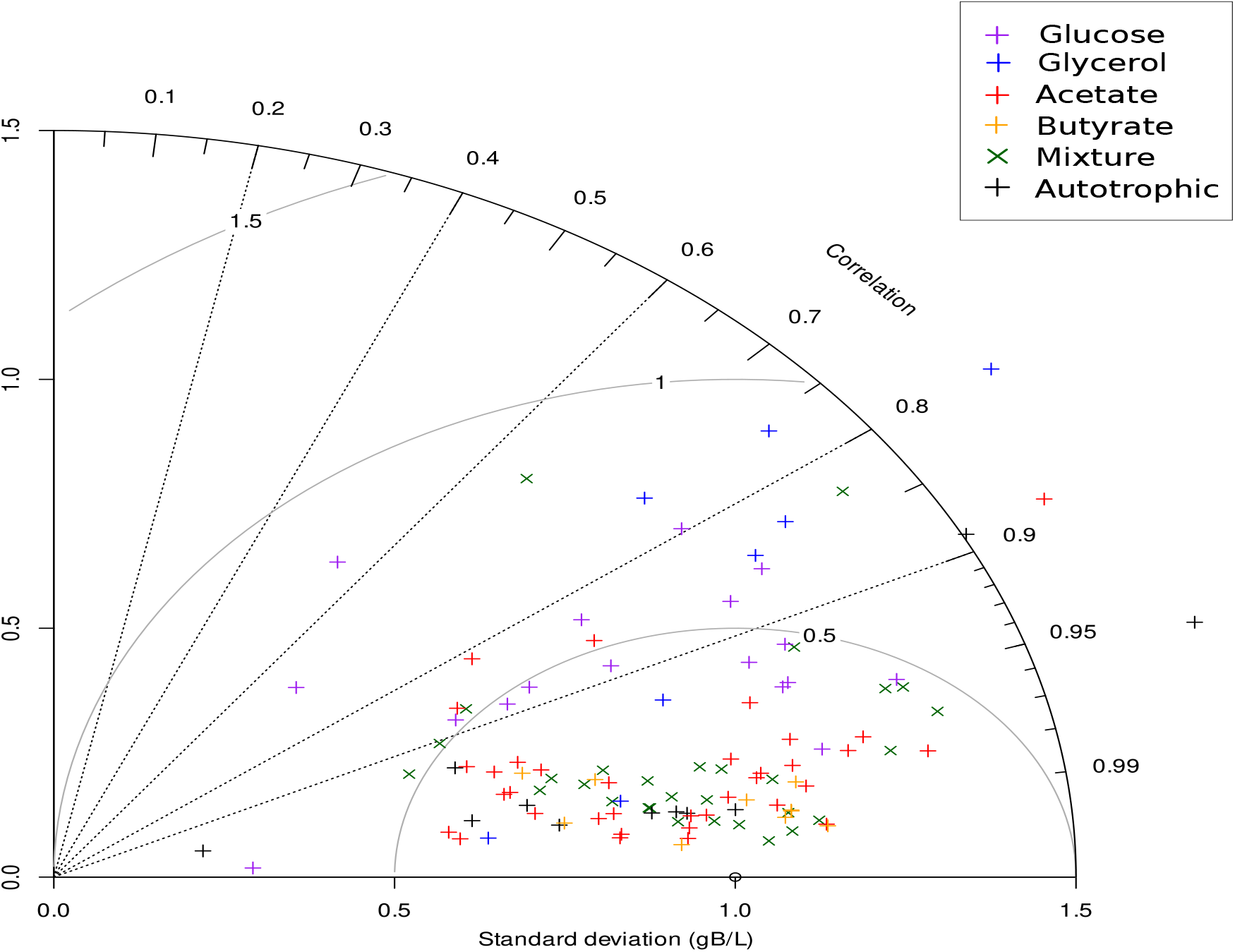
Taylor Diagram where each point represents the Pearson correlation coefficient and a normalized standard deviation of one experiment and model simulation. The semi circles centered at standard deviation 1.0 show the root-mean-square error.

**Figure 4:**
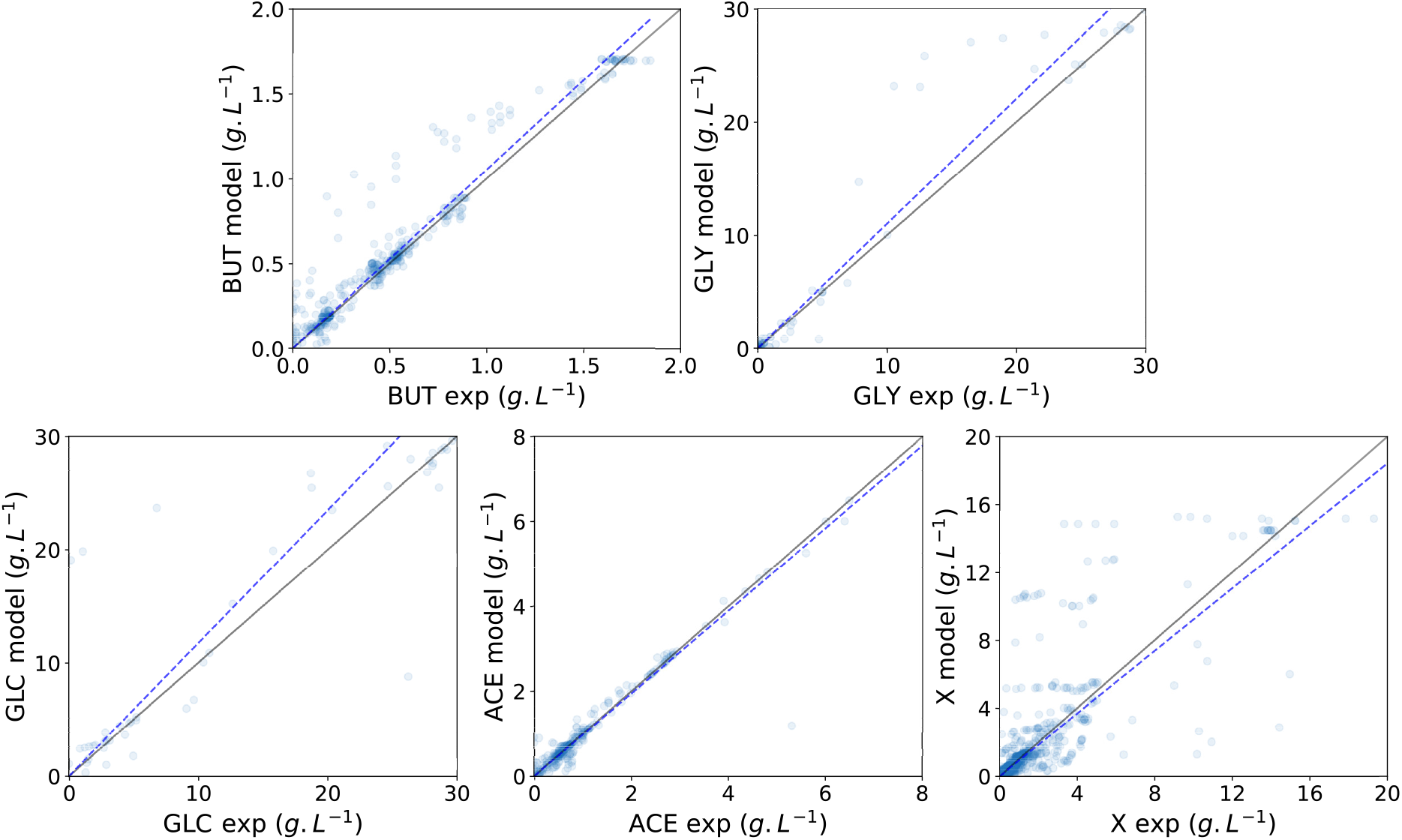
Validation of model predictions based on experimental data of butyrate, glycerol, glucose, acetate and biomass. All p-values for the regression are below 10^*−*3^. ℛ^2^ for the lines are, respectively, 0.97, 0.71, 0.95, 0.97 and 0.64. Darker colors represent a concentration of data points.

**Figure 5:**
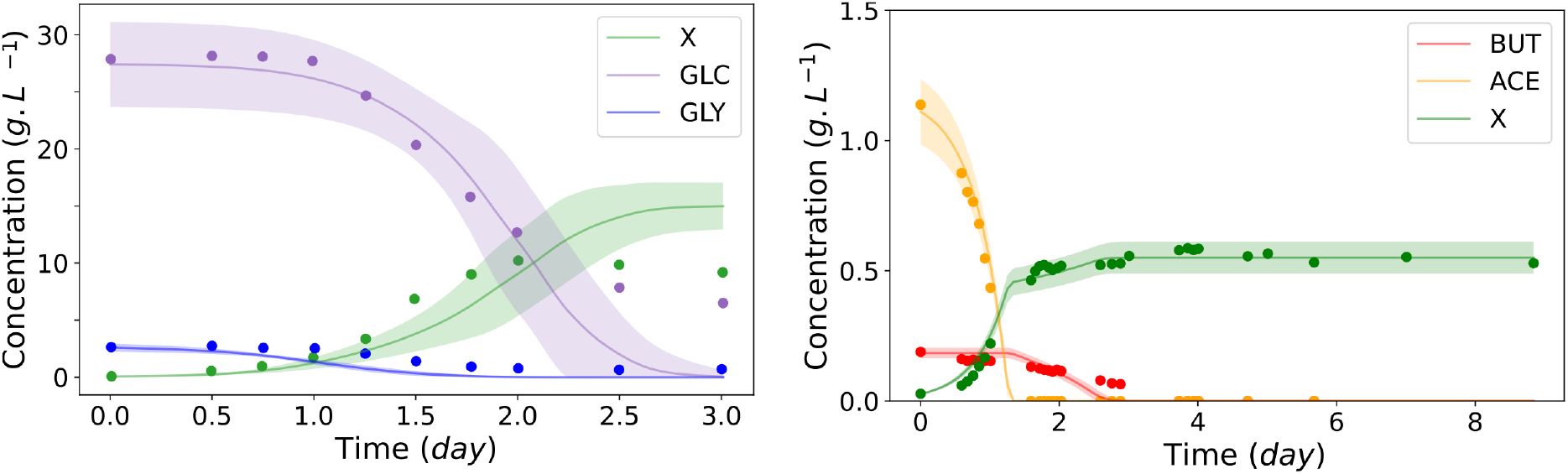
Model simulation and experimental data with two substrates present. From left to right: glycerol and glucose; butyrate and acetate. Colored regions represent ±1 standard deviation.

### 3.2. Analysis of metabolic maps

Thanks to the calculation of metabolic fluxes during the exponential growth phase, charts can be drawn to illustrate the metabolic fluxes within the cell (Figures 8 and 9). In this way, synthesis or degradation pathways can be predicted based on the presence or absence of one or several substrates, as well as the presence or absence of light.

During autotrophic growth, strong activity can be observed in the chloroplast subnetwork where photosynthesis occurs. More precisely, the greatest fluxes are associated with the fixation of *CO*_2_ by RUBISCO and the conversion to 3PG (3-phosphoglycerate) and the conversion of 3PG to glycerate-3-phosphate (GAP). GAP is then mainly transported toward the cytoplasm, where it is injected into the glycolysis. From then on, it enables the synthesis of precursor metabolites that are necessary for the synthesis of functional biomass composed of proteins, DNA, RNA, chlorophyll, carbohydrates, and lipids. During heterotrophic growth, no reaction occurs in the chloroplast subnetwork as no light is present. With a medium containing only acetate or butyrate, the largest fluxes are concentrated in the glyoxysome subnetwork where carbon substrates are converted into succinate. The synthesised succinate is exported from the glyoxysome and injected into the Krebs cycle. The latter then enables the synthesis of precursor metabolites for the production of functional biomass. To ascend the glycolysis, the anaplerotic pathways are in an upward direction.

No reaction takes place in the glyoxysome, neither in the glycerol utilization pathway for growth on glucose only. Glucose carried from the outside medium into the cytoplasm is directly injected into the upper glycolysis for the production of precursor metabolites necessary for the synthesis of functional biomass. Glycolysis is therefore entirely in a downward direction, and so are the anaplerotic pathways that enable the synthesis of oxaloacetate (OXA) that is needed to drive the Krebs cycle.

For growth on glycerol only, the greatest fluxes are located in the glycerol utilization pathway subnetwork, where imported glycerol in the cell is converted to glycerate-3-phosphate. The glycerate-3-phosphate is then injected into the middle of glycolysis, this time in an upward direction for the high glycolysis and in a downward direction for the lower glycolysis. The remnants of the fluxes are similar to those observed for glucose.

In mixotrophic conditions, when all carbon substrates are present in the medium, all metabolic reactions are activated. First, diauxic growth occurs, and acetate is consumed instead of butyrate. Glycerol and glucose are also used, but the flux remains after the acetate is depleted. Secondly, once all acetate has disappeared, butyrate is in turn consumed.

Figure 9 shows the metabolic fluxes after 30 hours of batch culture with optimal initial conditions (110 *g/l* of glucose and 20 *g/L* of glycerol). At this point, the acetate in the culture is practically depleted, but the concentration of butyrate remains similar to the initial condition. The rate of consumption of butyrate will remain stable for 2 days, and the rate will rapidly increase until the concentration threshold is reached. Because the optimal concentration is about 40 times lower than the threshold, the butyrate consumption rate does not reach the maximum value. Glycerol has a much lower concentration, while the flux of glucose concentration is still high. As a result of the higher concentration of biomass and the lower availability of light per cell, the metabolic fluxes in the chloroplast are reduced.

## 4. Discussion

### 4.1. Model perspectives

The reduced metabolic model we developed represents efficiently the microalgal growth under various substrates in heterotrophic or mixotrophic conditions. More accurate predictions could probably be obtained by complexifying the model to account for several factors. According to Lacroux et al. (2020), pH fluctuations when algae consume VFA can greatly impact growth. Associating a pH model that accounts for the various chemical species and their speciation, as proposed by Casagli et al. (2021a), would allow the calculation of the pH and the concentration of the undissociated form of the acids, which is the one actually taken up by the microalgae.

The effect of mixotrophic growth, here considered as the sum of autotrophic and heterotrophic conditions, is much more complex, and there is no consensus in the literature for a best-fit general model. For example, according to Martínez and Oruś (1991), mixotrophic growth is greater than the sum of autotrophic and heterotrophic conditions when the concentration of *CO*_2_ is limiting, since *CO*_2_ produced by fermentation can be recycled for the photosynthesis pathway. Nevertheless, under certain conditions, an inverse relationship between light intensity and glucose consumption can also occur (Yu et al., 2008, Patel et al., 2019, Wan et al., 2011).

Furthermore, the impacts of temperature, oxygen concentration, and *CO*_2_ concentration could result in considerable differences between the model and the experimental results. We also found that in some experiments with high concentrations of glucose and glycerol, another substrate was probably limiting the growth of the microalgae, most probably nitrogen. Although substrate inhibition is a well-known and important evolutionary mechanism (Reed et al., 2010), due to the above-mentioned issues, it is difficult to determine the exact optimal concentration of glucose and glycerol. Nevertheless, the usage of a model with inhibition results in better estimation of concentrations of substrates, while also accounting for other factors that might limit the microalgae growth rate in high concentrations of glucose and glycerol, e.g., oxygen limitation or low pH.

Extending the metabolic model to account for all these mechanisms is beyond the scope of this paper. It will require a large number of experiments to further calibrate and validate the model. Overall, despite the simplicity of the model in its present form, it is already very efficient. Especially when accounting for the diversity of strain and experimental conditions considered through the 15 studied papers. The model can then, already in its present form, can be used as a tool for optimizing microalgae growth on a mix of substrates.

### 4.2. Optimization of microalgal growth on inhibiting dark fermentation products

Acetate and butyrate are the main volatile fatty acids (VFA) products of dark fermentation. Butyrate will inherently lead to growth inhibition, and a strategy must be found to unblock this inhibition. Here, we sought, for batch and fedbatch cultivation strategies, to enhance microalgae growth in dark fermentation waste, by adding glucose and glycerol. We first studied mixotrophic conditions, with the effluent to treat considered as a typical dark fermentation product having concentrations of 3.5*g/L* of butyrate and 1.7*g/L* of acetate (Lacroux et al., 2020, Ghimire et al., 2015) and a continuous light intensity assumed at 500 *μmol/*(*m*^2^.*s*). The objective was that the chemical oxygen demand (COD) of the effluent at the end of the effluent treatment must be below 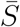 (here 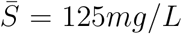) so that it can satisfy the state policies for discharge in the environment. We call this objective the regulation threshold.

This results in an optimal control problem (Harmand et al., 2019), whose objective is to minimize the time *t*_*f*_, where the COD of the remaining waste substrates falls below the regulation threshold. The control variable is the concentration of the substrate to be added to the dark fermentation effluent (glucose or glycerol).

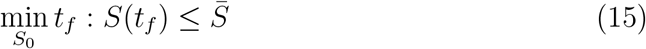

To solve the minimization problem we use a Nelder-Mead algorithm (Gao and Han, 2010). The output function simulates the metabolic model for a given *S*_0_ returning *t*_*f*_, the time required to reach the threshold. Considering that glucose can be added, it turns out that the addition 98*g/L* of glucose reduces the time to reach the COD threshold to 4.0 days compared to 16 days without the addition of carbon substrates (see Figure 6). Using only glycerol takes longer: 5.6 days with an addition of 39.5*g/L* glycerol. Using a mixture of both substrates, the optimal starting concentration for the batch is 115*g/L* of glucose and 19*g/L* of glycerol, reducing the time to consume the VFAs to only 2.7 days.

**Figure 6:**
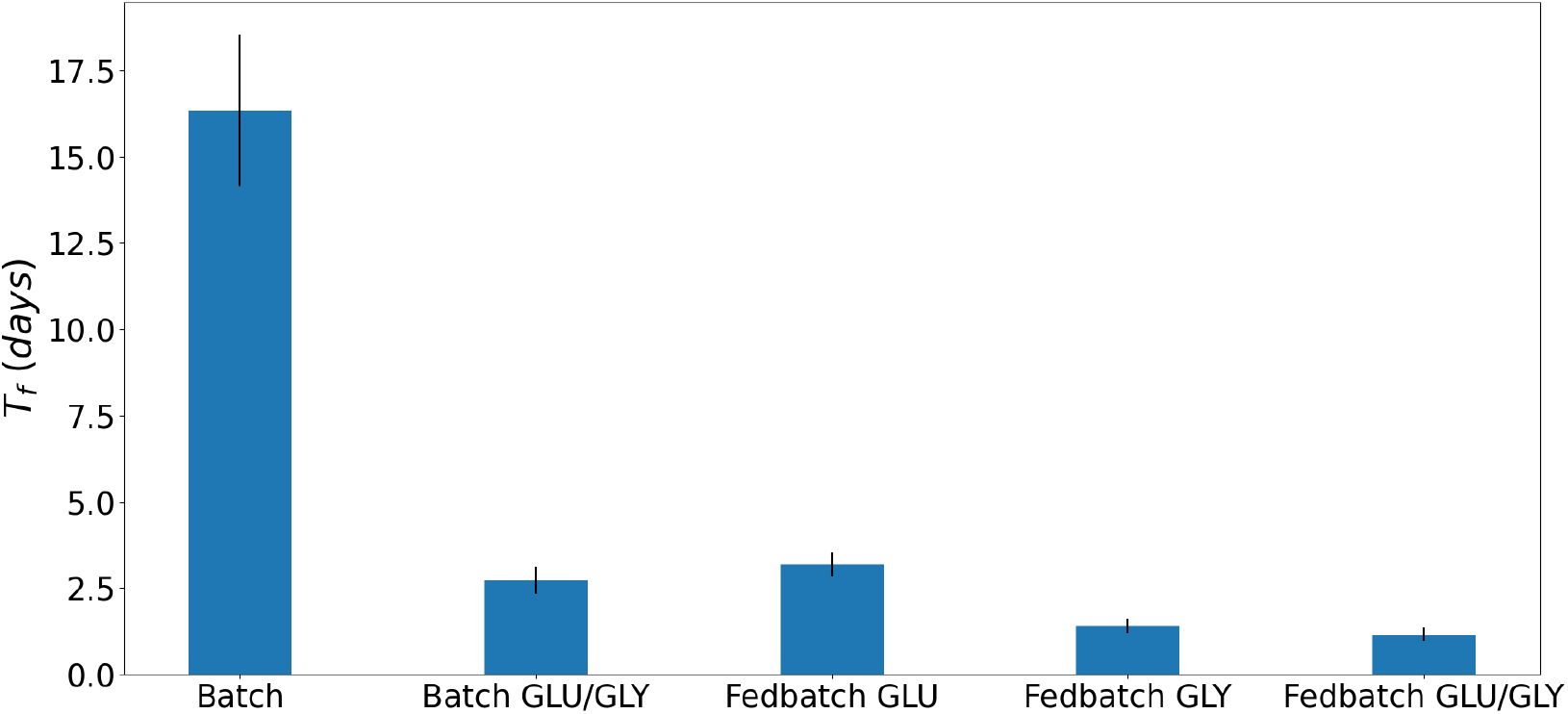
Time to reach the regulation threshold of a typical dark fermentation effluent using different conditions of cultivation - Batch (typical waste effluent), Batch GLC/GLY (typical waste with addition of an optimal concentration of glucose and glycerol), Fed-batch GLC, GLY, GLC/GLY (typical waste feeding, respectively, glucose, glycerol and a mixture of both.)

**Figure 7:**
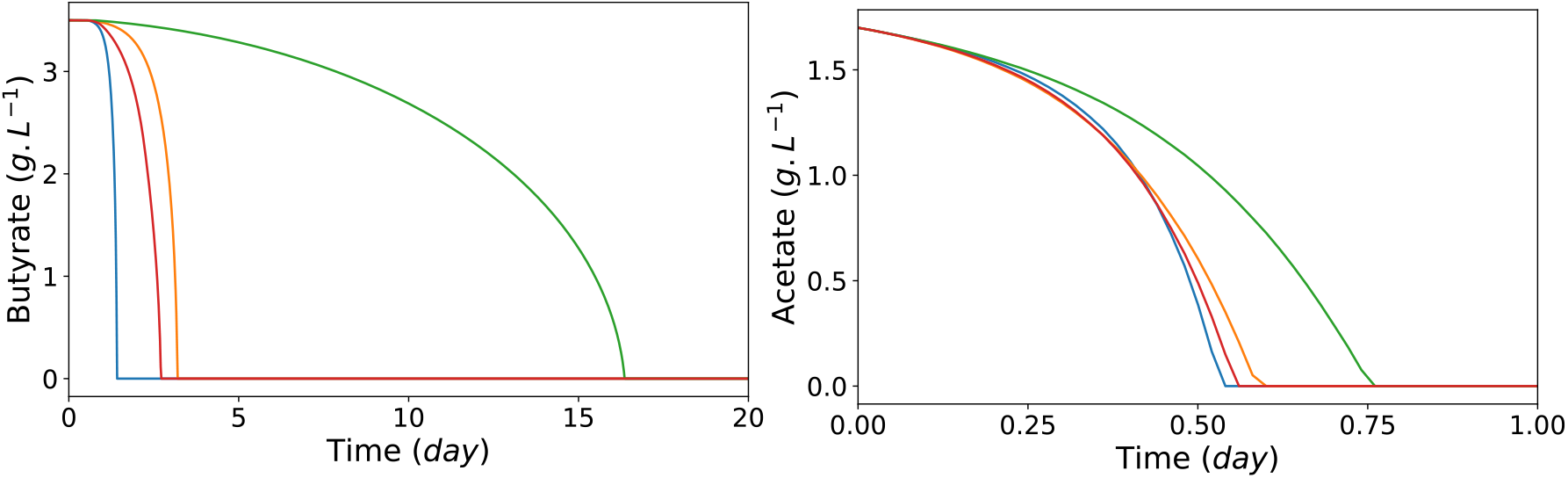
Concentrations of butyrate and acetate over time in different cultivation conditions. Optimal fed-batch with glycerol (blue), optimal fed-batch with glucose (red), batch with optimal addition of glycerol and glucose (orange), batch with only butyrate and acetate as substrates (green).

**Figure 8:**
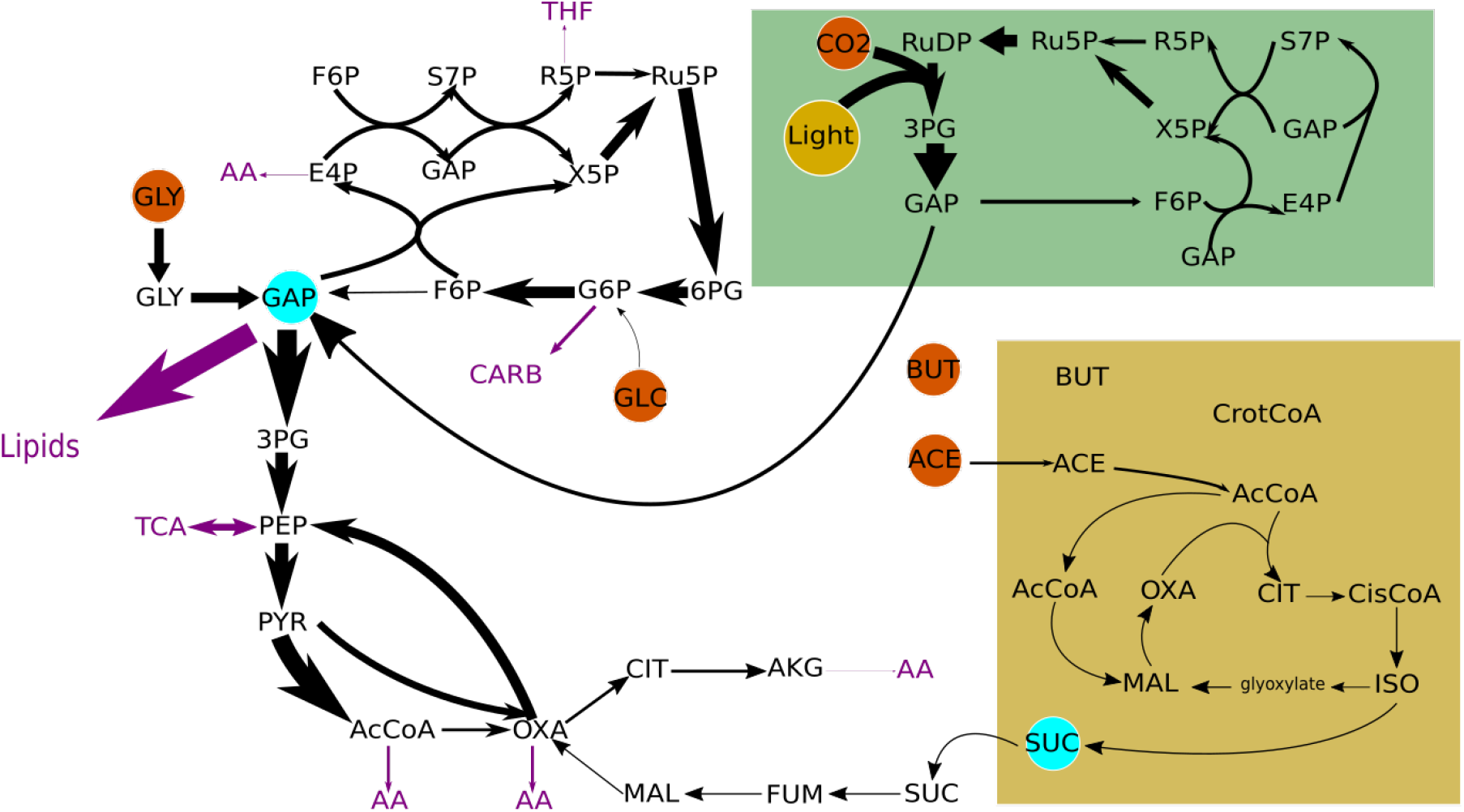
Metabolic chart showing the fluxes at initial concentration of acetate and butyrate with optimal conditions of glucose and glycerol.

**Figure 9:**
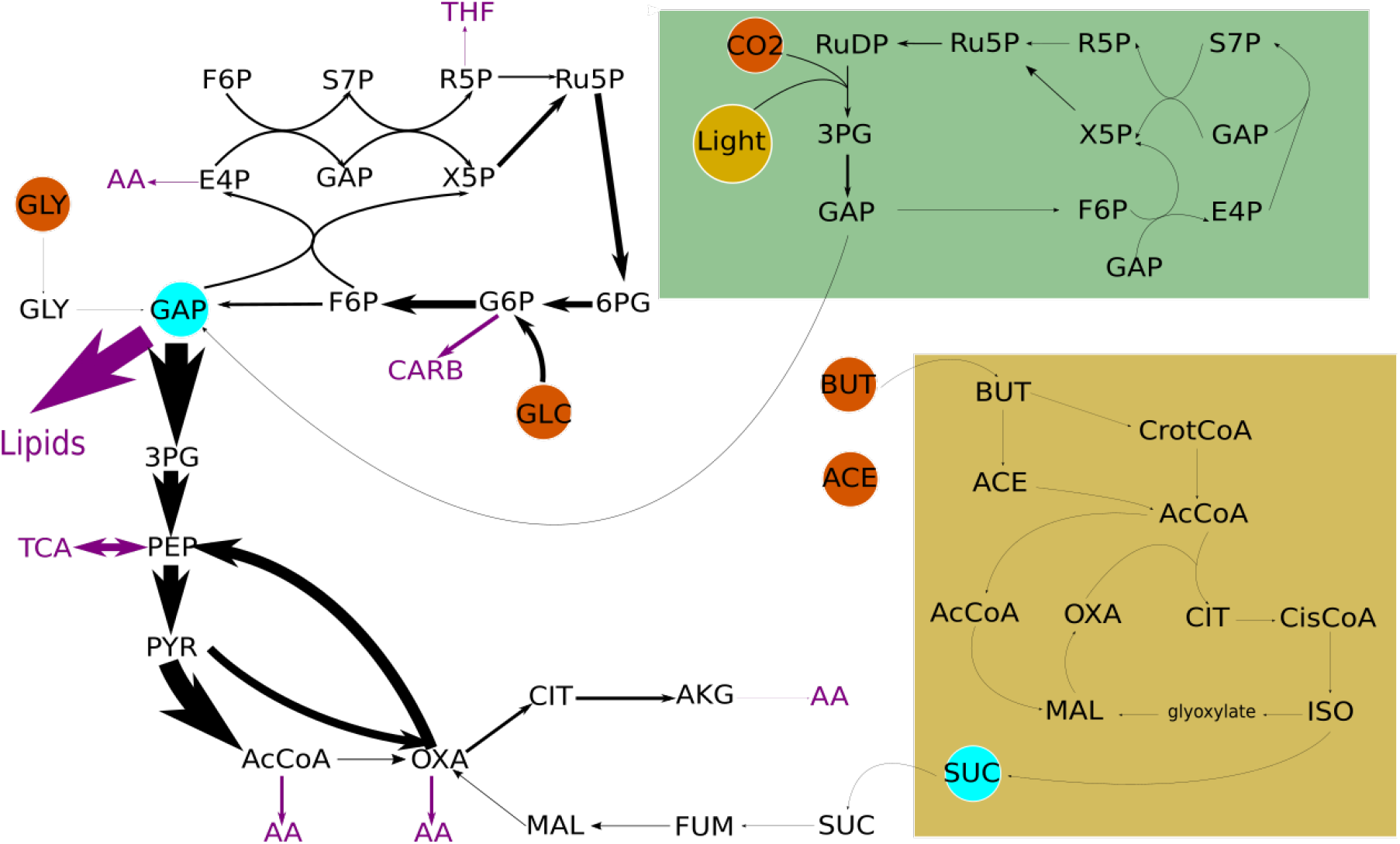
Metabolic chart showing fluxes for a batch process 30 hours after beginning of the batch.

Considering now a fed-batch cultivation systems instead of a batch one, the minimisation problem can be rewritten as following:

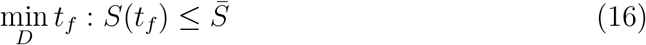

Where D is the dilution rate of the inflow containing only the additional substrates, at a high concentration, so that the volume of the reactor does not change. The optimal strategy can be approximated into a sub-optimal strategy, which would maximize the reaction rates for glucose and glycerol consumption. The strategy thus consists in computing the dilution rate such that glucose and glycerol concentrations stay constant close to the values for which the consumption of glucose and glycerol is maximum. The control problem is then reduced to finding the optimal stopping time of the inlet flux in the cases of a glucose, glycerol, and mixture inlet. In the case of the mixture, a fraction of 0.21 of glucose and 0.79 of glycerol is calculated, keeping glycerol at an optimal concentration. Using this control strategy, the final times for fed-batch cultivation are: 3.2 days for glucose, 1.4 days for glycerol and 1.2 days for a mixture of glucose and glycerol (see Figure 6).

Thanks to this approach, it will become possible to streamline two-stage water treatment strategies, and to recycle, carbon, nitrogen and phosphorus into the microalgal biomass.

## Acknowledgment

The authors acknowledge the support of the ControlAB ANR project (ANR-20-CE45-0014) and of the BIOMSA Ademe project. The UCAJEDI and EUR DS4H investments in the Future projects managed by the National Research Agency (ANR) with the reference numbers ANR-15-IDEX-0001.

